# Verbal semantic expertise is associated with reduced functional connectivity between left and right anterior temporal lobes

**DOI:** 10.1101/2024.04.11.589063

**Authors:** Wei Wu, Paul Hoffman

## Abstract

The left and right anterior temporal lobes (ATLs) encode semantic representations. They show graded hemispheric specialisation in function, with the left ATL contributing preferentially to verbal semantic processing. We investigated the underlying causes of this organisation, using resting-state functional connectivity as a measure of functional segregation between ATLs. We analysed two independent resting-state fMRI datasets (N=86 and N=642) in which participants’ verbal semantic expertise was measured using vocabulary tests. In both datasets, people with more advanced verbal semantic knowledge showed weaker functional connectivity between left and right ventral ATLs. This effect was highly specific. It was not observed for within-hemisphere connections between semantic regions (ventral ATL and inferior frontal gyrus; IFG, though it was found for left-right IFG connectivity in one dataset). Effects were not found for tasks probing semantic control, non-semantic cognition or face recognition. Our results suggest that hemispheric specialisation in the ATLs is not an innate property but rather emerges as people develop highly detailed verbal semantic representations. We propose that this effect is a consequence of the left ATL’s greater connectivity with left-lateralised written word recognition regions, which causes it to preferentially represent meaning for advanced vocabulary acquired primarily through reading.

## Introduction

Our understanding of the world is shaped by our semantic knowledge (Jefferies, 2013; Lambon Ralph et al., 2017). Convergent evidence from neuropsychology (Bozeat et al., 2000; Butler et al., 2009; Damasio et al., 2004), functional neuroimaging (Binder et al., 2011; Rice, Lambon Ralph, et al., 2015; Visser & Lambon Ralph, 2011) and brain stimulation studies (Lambon Ralph et al., 2009; Pobric et al., 2007, 2010) implicates both left and right anterior temporal lobes (ATLs) as critical regions for this semantic representation. One interpretation of these data could be that the semantic system shows redundancy, with both ATLs representing the same types of semantic information (Gainotti, 2012; Lambon Ralph et al., 2010; Schapiro et al., 2013; Snowden et al., 2012). However, there is limited empirical support for this *fully undifferentiated view*, since neuropsychological and neuroimaging studies suggest differential involvement of left and right ATL in different semantic tasks. Damage to left ATL has a greater impact on verbal semantic processing while right ATL damage can produce more significant deficits for faces and pictures (Butler et al., 2009; Gainotti, 2007, 2013; Rice et al., 2018; Snowden et al., 2012). Left ATL has also been particularly implicated in naming and speech production (Damasio et al., 2004; Hoffman & Lambon Ralph, 2018; Lambon Ralph et al., 2001; Rice, Lambon Ralph, et al., 2015; Woollams et al., 2017), while right ATL may be more associated with social semantic processing (Olson et al., 2007; Zahn et al., 2007). These findings have led some researchers to propose a *fully specialised view* (Gainotti, 2012, 2014; Snowden et al., 2004; Snowden et al., 2012) whereby left and right ATLs represent different forms of semantic knowledge associated with different modalities.

Between the extreme undifferentiated and fully specialised positions lies a *graded specialisation view* (Guo et al., 2013; Hoffman & Lambon Ralph, 2018; Rice, Hoffman, et al., 2015; Rice, Lambon Ralph, et al., 2015). Similar to the fully specialised view, the graded specialisation view argues for differences in the function of left and right vATLs. However, the graded specialisation model suggests that specialisation is relative rather than absolute, with a relative left ATL bias for tasks requiring verbal knowledge or verbal output. On this view, the ATLs together act as an integrative “hub” for semantic knowledge representation (Hoffman & Lambon Ralph, 2018; Lambon Ralph et al., 2010; Rice, Hoffman, et al., 2015; Rice, Lambon Ralph, et al., 2015). This graded specialisation is similar to graded preferences for word recognition and face recognition in left and right ventral occipitotemporal cortex (VOTC) (Cohen & Dehaene, 2004; Kanwisher et al., 1997; Puce et al., 1996; Thierry & Price, 2006). A similar pattern of graded specialisation has also been observed in inferior frontal gyrus (IFG), a key area for control and regulation of semantic processing (Badre & Wagner, 2007; Hoffman et al., 2010; Thompson-Schill et al., 1997; Vitello & Rodd, 2015). Parts of IFG show a left-hemisphere bias for verbal semantic tasks and a right-hemisphere bias for non-verbal tasks (Krieger-Redwood et al., 2015). Thus, graded hemispheric specialisation appears to be a feature of semantic processing across multiple brain regions.

Though there is now considerable evidence for graded hemispheric specialisation across the ATLs, little attention has been paid to how to this specialisation develops. In other neural systems, the development of specific cognitive abilities drives increasing functional specialisation (Guerra-Carrillo et al., 2014; Uddin et al., 2010). In VOTC, for example, right-lateralised responses to faces emerge as children learn to read (Dehaene et al., 2015; Monzalvo et al., 2012) and the strength of this lateralisation is correlated with reading competence (Dehaene-Lambertz et al., 2018; Dundas et al., 2013). One explanation for this phenomenon, supported by computational models, is that orthographic processing is tightly coupled with left-lateralised speech production systems that are anatomically connected with left but not right VOTC (Behrmann & Plaut, 2020; Plaut & Behrmann, 2011). This asymmetry causes neurons in left VOTC to specialise for word recognition and, consequently, more of the computational work for face processing is taken up by the right VOTC. The result is increasing hemispheric specialisation within a generally bilateral object recognition system. A similar mechanism could be at play in the ATLs (Hoffman & Lambon Ralph, 2018; Woollams & Patterson, 2018). From later childhood onwards, most advanced vocabulary knowledge is acquired through exposure to print (Cunningham & Stanovich, 1998). The development of advanced verbal semantics is therefore closely linked with the left-lateralised process of written word recognition, and this connectivity bias could lead the left ATL to specialise for representing verbal semantics. Right ATL specialisation for non-verbal aspects of knowledge might then be an emergent consequence.

In the present study, we used resting-state fMRI to test the hypothesis that greater verbal semantic expertise is associated with greater hemispheric specialisation in the semantic system. Resting-state functional connectivity (RSFC) is commonly used to measure the development of functional neural networks. In childhood, increasing age is associated with greater segregation (i.e., weaker correlations) between distinct networks, suggesting increasing differentiation in function (He et al., 2019; Uddin et al., 2010). For example, functional connectivity between left and right VOTC is lower in adults than in children (He et al., 2015). In adulthood, the acquisition of new skills is also associated with changes in the resting-state connectivity patterns of relevant networks (Guerra-Carrillo et al., 2014; Zhu et al., 2011). Acquisition of advanced semantic knowledge might therefore drive functional specialisation that can be observed through RSFC. There is already some evidence for this at a global level: Wang et al. (2021) found that young adults with greater crystalised intelligence displayed greater segregation between large-scale brain networks. Here, we took a more targeted approach to test whether the same is true for core nodes of the semantic system in ATL and IFG.

Our hypothesis is that the development of verbal semantic expertise in adulthood is associated with increasing specialisation in function between left and right ATLs, as the left ATL comes to disproportionately represent verbal knowledge acquired through reading. There are two ways that this functional specialisation might manifest itself. First, older people might show greater segregation between left and right ATLs than young people because they have accumulated more verbal semantic knowledge during their lives (Grady, 2012; Hoffman, 2018, 2019; Park et al., 2002; Verhaeghen, 2003; Wu & Hoffman, 2022, 2023a). Second, independent of age, people with more extensive verbal knowledge (indexed by vocabulary tests) may show greater segregation between ATLs. We tested these possibilities in two independent datasets that include RSFC data from a range of age groups. We focused on connectivity between the ventral surfaces of the ATLs (vATLs) as this region shows the most consistent involvement in semantic processing across a range of categories and input modalities (Lambon Ralph et al., 2017).

We also investigated whether parallel effects occur in the IFGs. IFG is involved in regulation and control of semantic processing, rather than representation of knowledge per se (Jefferies, 2013; Lambon Ralph et al., 2017). Less is known about how these functions are arranged across left and right IFGs but there is some evidence for graded specialisation mirroring that seen in the ATLs (Krieger-Redwood et al., 2015). If interaction with written materials spurs the development of more specialised semantic control functions, then we would expect greater ability in verbal semantic control to be associated with more between-IFG segregation. Effects of ageing may differ here, however. Semantic control ability begins to deteriorate with age and older people appear to rely more on bilateral IFG activation to regulate their semantic processing (Hoffman, 2018, 2019; Hoffman & Morcom, 2018; Wu & Hoffman, 2022). Thus, segregation between IFGs might decrease with age.

## Method

We report two parallel sets of analyses on different datasets. We first conducted analyses on resting-state fMRI data collected as part of a larger study of age-related effects on semantic cognition (Wu & Hoffman, 2023a, 2023b). These data are publicly available (https://osf.io/zbxt4) and the resting-state fMRI data have not been reported previously. To replicate our findings, we then conducted validation analyses using data from the Cam-CAN project repository (available at http://www.mrc-cbu.cam.ac.uk/datasets/camcan) (Shafto et al., 2014; Taylor et al., 2017).

### Participants

Forty-five older adults and 45 young adults were recruited from the University of Edinburgh Psychology department’s volunteer panel and local advertising and participated in the study in exchange for payment. All participants were native English speakers and reported to be in good health with no history of neurologic or psychiatric illness. The older participants completed the Mini-Addenbrooke’s Cognitive Examination (Hsieh et al., 2015) as a general cognitive screen. We excluded data from two older participants, who scored < 26 of 30 on the Mini-Addenbrooke’s Cognitive Examination. Two young participants’ data were also excluded because of technical issues or structural abnormalities. Thus, data from 43 older participants (28 females, 15 males; mean age = 68.14 years, s.d. = 5.21 years, range = 60 - 79) and 43 young participants (31 females, 12 males; mean age = 23.07 years, s.d. = 3.23 years, range = 18 - 32) were used in the analyses. Both age groups had a high level of formal education (older adults years of education: mean = 15.65 years, s.d. = 2.84 years, range = 10 - 22; young adults: mean = 17.07 years, s.d. = 2.53 years, range = 12 - 23), and young adults had completed more years of education than older adults (*t* _84_ = 2.44, two-tailed *p* < 0.05). Informed consent was obtained from all participants and the research was performed in accordance with all relevant guidelines and regulations. The study was approved by the University of Edinburgh Psychology Research Ethics Committee. Sample size was determined by the resources available to complete the study.

#### Validation analyses

We used data from the Cam-CAN dataset (Shafto et al., 2014; Taylor et al., 2017) to conduct validation analyses. In this cohort, we first removed data from seven participants who had missing performance scores in more than eight behavioural tests across the two stages of the Cam-CAN project. This was because a large proportion of missing behavioural data may indicate the participant had abnormal cognitive ability. Then we focused on individuals whose fMRI data from resting-state and movie watching scans were both available. Data from 642 participants (324 females, 318 males; mean age = 54.65 years, s.d. = 18.46 years, range = 18.50 - 88.92) were used for analysis.

### Tasks

In our study, participants completed a verbal synonym judgement task, a verbal feature matching task and a cognitively demanding non-semantic task during fMRI. Neuroimaging data for these tasks have been reported elsewhere (Wu & Hoffman, 2023a, 2023b) and were not used in the current study. Here, we used participants’ behavioural responses during scanning as measures of semantic and general cognitive ability. The stimuli for all three tests were taken from the norms of Wu and Hoffman (2022) (see Figure 1 for examples).

**Figure 1.**
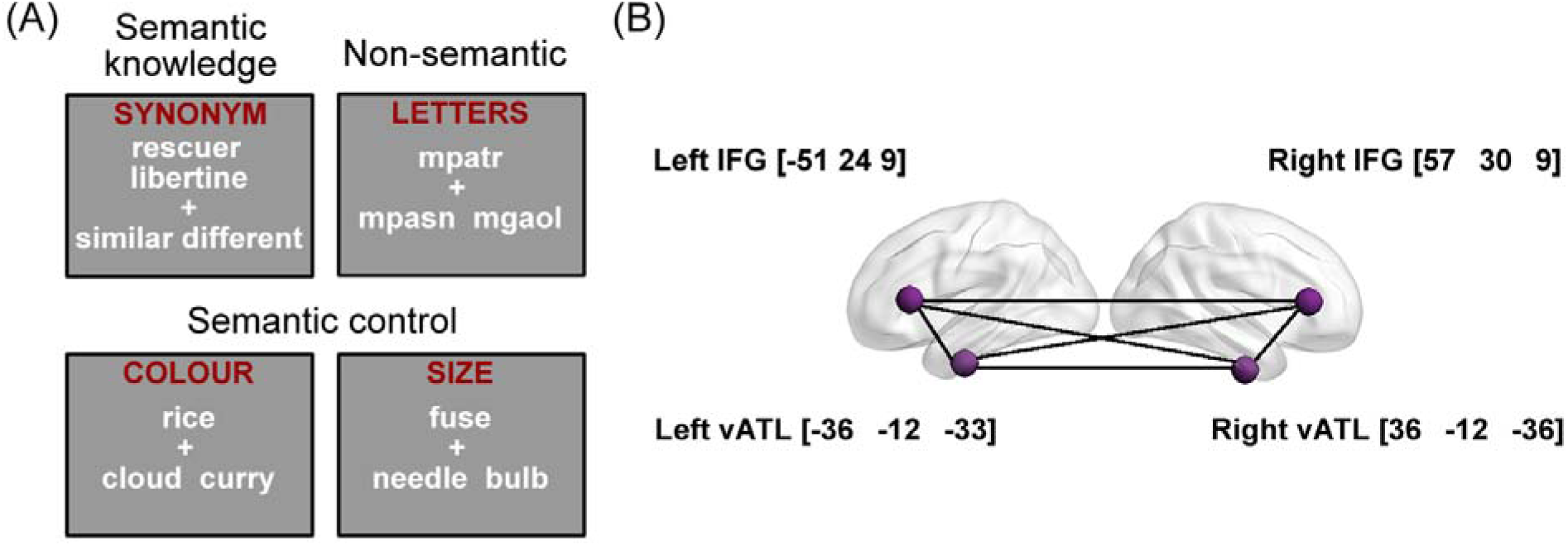
(A) Example items from each task in our dataset and (B) seed ROIs of left and right vATLs and IFGs and their MNI coordinates.

#### Test of verbal semantic knowledge

We used an 80-item synonym-judgment task to probe participants’ verbal semantic knowledge (i.e., vocabulary). On each trial, two words along with 2 options (i.e., similar or different) were shown to the participant around the centre of the screen. Participants were asked to decide if the two words shared a similar meaning or not. Because it used a high proportion of low-frequency words, whose meanings are less familiar to people, this task measured the breadth of verbal semantic knowledge that participants possessed.

#### Test of verbal semantic control

An 80-item feature-matching task was designed to probe participants’ ability to exercise cognitive control over the retrieval and selection of semantic knowledge. On each trial, participants were presented with a probe word along with 2 option words. They were asked to select the option that shared a particular semantic feature with the probe (40 colour trials and 40 size trials). For instance, on a colour trial, *sunflower* would match with *lemon* as both are typically yellow. This task demands semantic control processes as participants need to select the target semantic feature and inhibit competing but irrelevant semantic information from the distractor (e.g., *sunflower* - *stalk*) (Badre et al., 2005; Thompson-Schill et al., 1997). This was a verbal task as all stimuli were written words. However, because the features we used are related to visual properties of objects, it also required access to non-verbal aspects of semantic knowledge.

#### Test of non-semantic cognitive control

We used an 80-item string-matching task to examine participants’ general executive control ability. Each trial in this task consisted of 3 meaningless letter strings (e.g., *mpatr*), and participants were asked to choose which of the two options had greatest orthographic similarity with the probe (i.e., sharing the most letters in the same order). This task examined general cognitive control processes, as participants needed to resolve competition between options without engaging semantic processing. We included this measure in analyses to control for non-specific effects of cognitive performance.

#### Validation analyses

Performance data from two cognitive tests in the Cam-CAN battery were used: the Spot The Word test (STW, Baddeley et al., 1993) and a familiar face recognition test (Bartlett & Leslie, 1986). The STW test requires a participant to identify the real words out of pairs of items comprising one word and one non-word. This test was conducted in stage 1 of the Cam-CAN project as a measure of verbal intelligence (Taylor et al., 2017), and it is comparable to the semantic knowledge task in our dataset as both examined breadth of verbal knowledge (i.e., people with more advanced verbal knowledge should correctly recognise more words). We included the familiar face recognition test as a control. In this test, participants were shown 40 images of faces (30 famous and 10 unfamiliar) and asked to judge whether the face is familiar or not. For a face identified as familiar, participants were also asked to verbally provide the person’s name, and other information (e.g., occupation). The familiar face recognition task is a multimodal semantic test as it entails the linking of non-verbal (i.e., the face) with verbal semantic knowledge (i.e., the name). As such, we would not expect greater segregation between left and right vATLs to be associated with performance.

### Design and procedure

In our dataset, there were two task-fMRI scanning runs, in which participants completed the synonym judgement, feature matching and non-semantic tasks in a blocked fashion. Behavioural data from these two runs were used as performance measures in the current study. The neuroimaging data from these two runs were used to define the seed regions for resting-state analyses (see below). In between the two task-fMRI runs, resting-state fMRI data were acquired while participants rested with their eyes open. There was a fixation cross on the screen during the scan and this run lasted 6 minutes.

#### Validation analyses

We used data from two fMRI scans in the Cam-CAN data, including a resting-state scan which lasted for 8 min and 40 s, during which participants rested with their eyes closed and a movie-watching scan, in which participants passively viewed an 8-min excerpt of a film: Alfred Hitchcock’s “Bang! You’re Dead”. We included the movie data because the resting-state scan used a single-echo imaging sequence that is relatively insensitive to activation in the vATLs (Ojemann et al., 1997; Visser et al., 2010). In contrast, the movie scan used a multi-echo sequence that has been shown to improve signal quality in this brain region (see next section). By combining data from both scans, we therefore improved sensitivity to effects in vATLs. However, qualitatively similar results were obtained when analysing only the resting-state scan.

### Image acquisition and processing

For our dataset, images were acquired on a 3T Siemens Skyra scanner with a 32-channel head coil. fMRI in the vATL region is affected by susceptibility artefacts that can negatively impact data quality (Devlin et al., 2000). To combat this, we adopted a whole-brain multi-echo acquisition protocol shown to improve ventral temporal fMRI signal as well as minimising the effects of head movement (Halai et al., 2015; Kundu et al., 2017). The fMRI data were simultaneously acquired at 3 echo times (TEs), and then were weighted and combined and the resulting time series were denoised using independent component analysis (ICA). The multi-echo Echo-Planar Imaging (EPI) sequence included 46 slices covering the whole brain with TE at 13 ms, 31 ms, and 50 ms, repetition time (TR) = 1.7 s, flip angle = 73°, 80 × 80 matrix, reconstructed in-plane resolution = 3 mm × 3 mm, slice thickness = 3.0 mm, and multiband factor = 2. One run of 212 volumes was acquired. For each participant, a high-resolution T1-weighted structural image was also acquired using an MP-RAGE sequence with 1 mm isotropic voxels, TR = 2.5 s, TE = 4.4 ms.

Images were preprocessed and analyzed using SPM12, the TE-Dependent Analysis Toolbox (Tedana) (DuPre et al., 2021) and the DPABI Toolbox (Yan et al., 2016). The first 4 volumes of the time series were discarded. Estimates of head motion were obtained using the first BOLD echo series. Slice-timing correction was carried out and images were then realigned using the previously obtained motion estimates. Tedana was used to combine the 3 echo series into a single-time series and to divide the data into components classified as either BOLD-signal or noise-related based on their patterns of signal decay over increasing TEs (Kundu et al., 2017). Components classified as noise were discarded. After that, images were unwarped with a B0 fieldmap to correct for irregularities in the scanner’s magnetic field. Functional images were spatially normalized to MNI space using SPM’s DARTEL tool (Ashburner, 2007). To improve the quality of the resting-state fMRI signals, additional preprocessing procedures were conducted via DPABI (https://rfmri.org/DPABI, Yan et al., 2016), including: (1) linear detrending, (2) regression of motion parameters (Friston-24 parameters) (Friston et al., 1996), mean white matter (WM) and cerebrospinal fluid (CSF) signals, global signal, as well as outlier scans which had a large framewise displacement (FD), (3) temporal band-pass (0.01–0.1 Hz) filtering, and (4) spatial smoothing with a Gaussian kernel of 6 mm full width half maximum (FWHM). We chose to remove the whole-brain signal in the second step to reduce the effect of physiological artifacts (Birn et al., 2006; Fox et al., 2005; Yan & Zang, 2010).

#### Validation analyses

We used preprocessed fMRI images from the Cam-CAN dataset. The resting-state fMRI data had 261 volumes and acquired with 32 slices with 3.7 mm slice thickness, TR = 1.97 s, TE = 30 ms, flip angle = 78°, FOV = 192 mm × 192 mm, resolution = 3 mm × 3 mm × 4.44 mm. As described by Taylor et al. (2017), preprocessing involved unwarping, realignment, slice-time correction and normalisation. Data-driven wavelet despiking was applied to minimize motion artefacts (Patel et al., 2014). For the movie-watching data, there were 193 volumes scanned with a multi-echo EPI sequence, TR = 2.47 s, TE = 9.4 ms, 21.2 ms, 33 ms, 45 ms and 57 ms, flip angle = 78°, FOV = 192 mm × 192 mm, resolution = 3 mm × 3 mm × 4.44 mm. The preprocessing procedures for the movie-watching data were similar to the resting-state data, except that ICA was performed to combine the 5 echo series. For both the resting-state and movie-watching data, we used DPABI to perform the same set of additional preprocessing procedures (with the same parameters) as for our dataset, including linear detrending, regression of nuisance variables, band-pass filtering and smoothing. Note that, because the Cam-CAN dataset had already undergone wavelet despiking processing, a new data-driven method that can be used to control head movement artifacts and replace traditional data scrubbing (Patel et al., 2014), we did not regress out outlier scans which had a large FD for the Cam-CAN dataset.

### Definition of regions of interest (ROIs)

To identify seed regions in left and right vATLs and IFGs, we used a method that combined anatomic masks with group-level task-activated peaks. Several steps were involved in this method.

First, we generated anatomical vATL and IFG masks. We defined the IFGs using the BA 45 mask in the Brodmann Areas Map provided with MRIcron (https://people.cas.sc.edu/rorden/mricro/template.html). We defined anatomical vATL regions in a similar fashion to Hoffman and Lambon Ralph (2018). That is, we generated an anatomical mask using the voxels with a greater than 50% probability of falling within fusiform gyrus in the LONI Probabilistic Brain Atlas (LPBA40), and we divided the fusiform gyrus mask into 5 roughly-equal-length sections that ran along an anterior-to-posterior axis. The anatomical masks of vATLs were then constructed by combining the anterior two sections of the above mask.

Second, using the two task-fMRI runs in our dataset (see Design and procedure section), we contrasted the average of the synonym judgement and feature matching tasks versus the non-semantic task to identify voxels that were most responsive to semantic processing. By overlapping this functional map with the anatomical masks, we obtained group-level peak coordinates in the left and right vATLs and IFGs. Spheres with 8-mm radius were built centred on these coordinates, which were the final seed ROIs used in the functional connectivity analyses (shown in Figure 1).

### Resting-state functional connectivity (RSFC) analyses

The same set of analyses were conducted for both our dataset and the Cam-CAN dataset. For each participant, timecourse in each seed ROI (i.e., left and right vATLs and IFGs) was extracted by averaging across all voxels in the ROI. Pearson correlations between timecourses were calculated for each pair of seeds. The resulting correlation coefficients were then Fisher-transformed to Z scores, which were used for group-level analyses. For the Cam-CAN data, this process was performed separately for the resting-state and movie-watching scans and the obtained Z scores were averaged for each participant to give a single Z score.

We first examined how intrinsic functional connectivity within the core semantic network (i.e., vATLs and IFGs) varied with age. For our dataset, linear regression models were built to predict RSFC values of each pair of seed ROIs using age group as a predictor. To account for potential effects of head motion on RSFC estimates, we also calculated the individual-level mean framewise displacement (mean FD) (Power et al., 2012) during the resting-state scanning and included it in each model as a covariate of no interest.

#### Validation analyses

For the Cam-CAN dataset, which includes participants across a wide range of ages, we treated age as a continuous variable and used it to predict the RSFC values of each pair of seed ROIs (i.e., the Fisher-z-transformed correlation values). The individual-level mean FD in the resting-state and movie-watching scans were averaged and included in the models as a covariate of no interest.

We next constructed a series of further linear regression models to investigate the relationship between RSFC and participants’ performance in the cognitive tasks. For each task in our dataset, we specified a regression model with age group, task performance (i.e., accuracy) and their interaction as predictors of RSFC values for each pair of seeds. Mean FD was included in the model as a covariate of no interest. This model was then compared with the previous model (i.e., the age effect model) to examine if the inclusion of task performance improved model performance in predicting RSFC values. These models revealed how task performance and its interaction with age were correlated with RSFC for each pair of ROIs. All continuous predictors were standardized prior to entry in our models. We used accuracy rather than reaction time as the behavioural measure because: (1) reaction times vary between age groups due to changes in general processing speed that are not specific to semantic cognition (Salthouse, 1996), and (2) we aimed to keep our behavioural measures consistent with the behavioural measures used in the Cam-CAN data.

#### Validation analyses

We used the total score in the STW task and performance in the face recognition task (i.e., number of faces correctly recognised as familiar subtracting false alarms) as the behavioural measures for the Cam-CAN dataset. For each task, we built a regression model with age and task performance as predictors to predict the RSFC values of each pair of seeds, and individual-level mean FD was included in the model as a covariate of no interest. This model was then compared with the corresponding age-only model. Age was treated as a continuous variable in the Cam-CAN models. All regression models were computed using the lme4 package in R (https://cran.r-project.org/) and p-values were corrected for multiple comparisons using the false discovery rate approach (Benjamini & Hochberg, 1995).

It is worth noting that, although the RSFC of left-right IFGs and left-right vATLs were the main focus of the current study, we also investigated the other connections between IFGs and vATLs for completeness. We report the two key cross-hemispheric connections in the main paper and show the effects for other connections in Supplementary Information.

## Results

Using our dataset, we began by testing how connectivity between the left and right vATLs and IFGs varied across age groups. When controlling for FD, older people showed significantly stronger correlations between IFGs than young people (see Table 1 and Figure 2). They showed weaker correlations between vATLs than young people but this effect did not survive correction for multiple comparisons (*p* = 0.07). Thus there was only weak support for the idea that functional specialisation in the vATLs increases with age. Other connections did not vary between age groups (see Supplementary Table S1 and Figure S1). We then tested whether cross-hemispheric RSFC was related to performance in semantic and non-semantic tasks. We did this by adding performance on each of the three tasks (and their interaction with age) separately to the model in turn and testing whether this improved model fit. The addition of semantic knowledge scores improved the fit of the left vATL - right vATL (*F* = 7.53, *p* < 0.01) and left IFG - right IFG (*F* = 4.73, *p* < 0.05) models. As shown in Table 2 and Figure 3, people with better performance on the synonym judgment task exhibited weaker left vATL - right vATL connectivity (performance effect: *B* = -0.094, *SE* = 0.028, *p* < 0.01) and weaker left IFG - right IFG connectivity (performance effect: *B* = -0.117, *SE* = 0.040, *p* <0.01). Thus, people with greater verbal semantic expertise appeared to show more functional specialisation between left and right semantic regions. For the vATLs this was qualified by an age × performance interaction, as the semantic knowledge effect was stronger in older people. Effects of age remained significant for IFGs but not vATLs (see Table 2), suggesting that the weak effect of lower bilateral vATL connectivity in later life could be due to development of more advanced verbal semantic knowledge. Semantic knowledge did not predict the strength of other IFG-vATL connections (i.e., adding these predictors did not improve model fit, *F*s <= 0.49, *ps* >= 0.61; see Supplementary Table S2 and Figure S2 for results of these models). Feature-matching task performance and non-semantic task performance were not associated with connection strength for any connections, indicating that effects were specific to the verbal knowledge representation and not to other aspects of semantic cognition or general cognitive ability.

**Figure 2.**
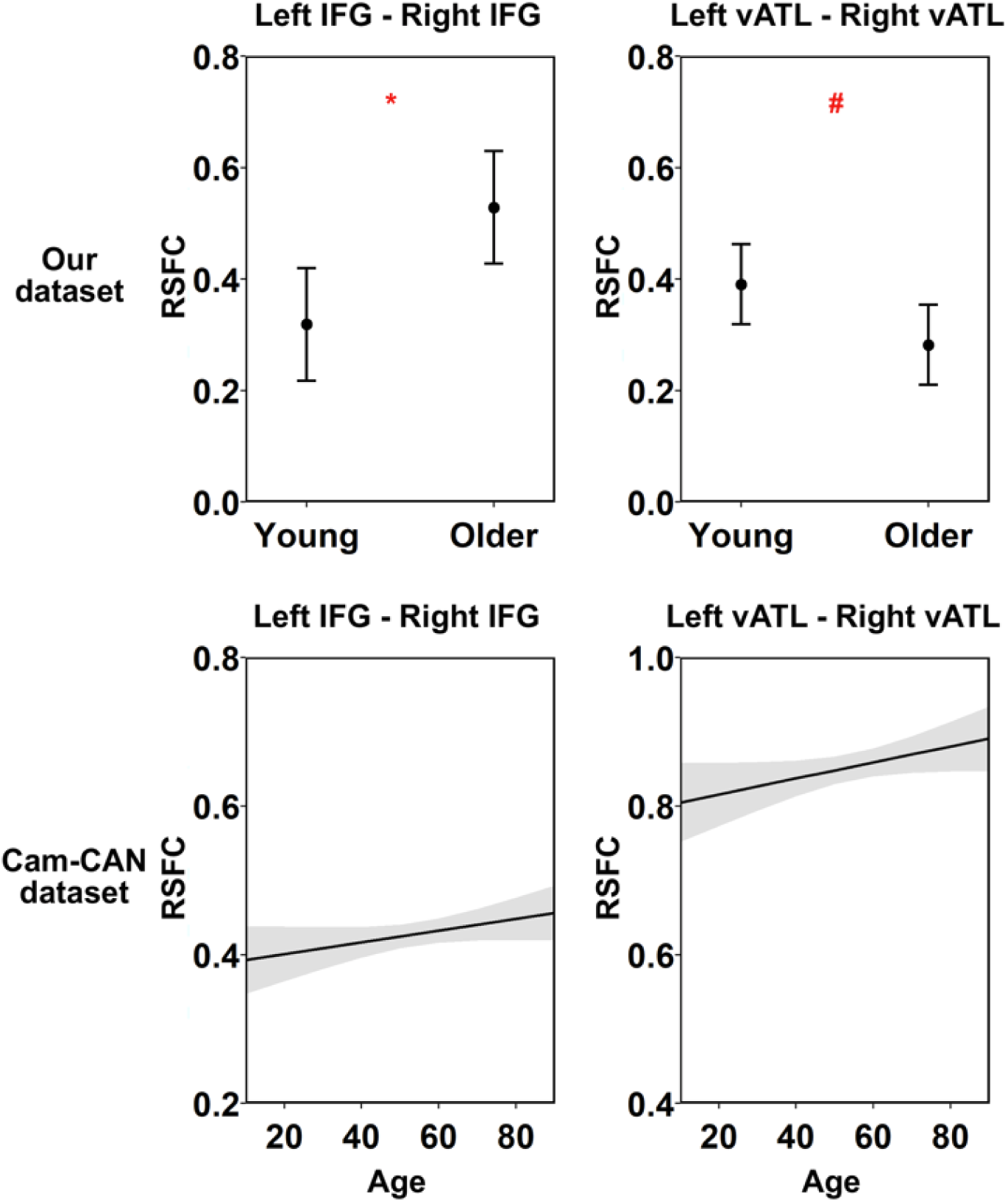
Results of linear regression models for age effects. This figure shows the modelled effects of age on RSFC between each pair of seed ROIs in each task. Shadow areas and error bars indicate 95% confidence intervals. The asterisks indicate significance level after FDR correction within dataset, # *p* = 0.070, * *p* < 0.05, ** *p* < 0.01, *** *p* < 0.001.

**Figure 3.**
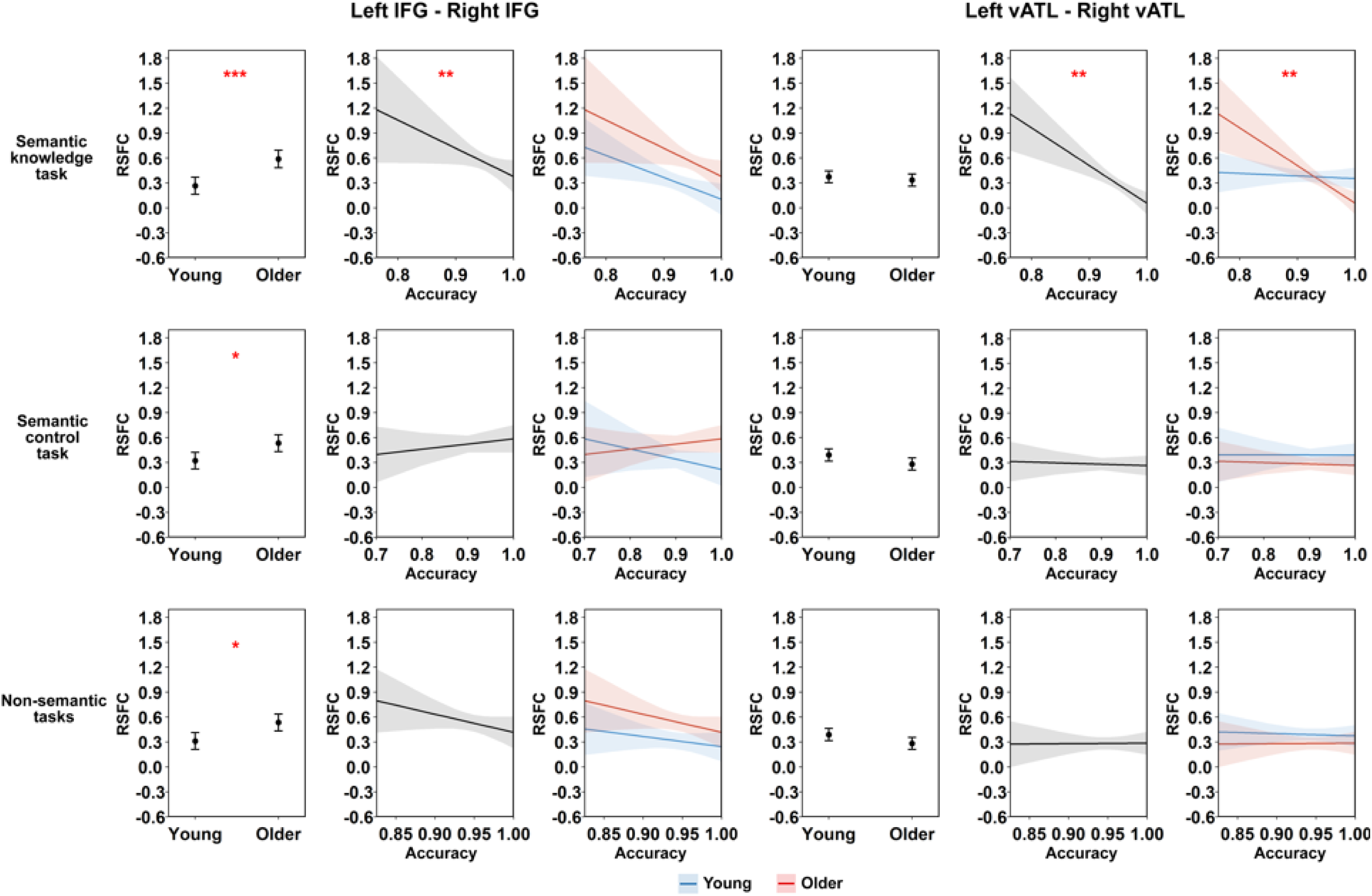
Results of linear regression models for our dataset. This figure shows the modelled effects of age group and task performance on RSFC between each pair of seed ROIs. Shadow areas and error bars indicate 95% confidence intervals. The asterisks indicate significance level after FDR correction within each task, * *p* < 0.05, ** *p* < 0.01, *** *p* < 0.001.

**Table 1.**
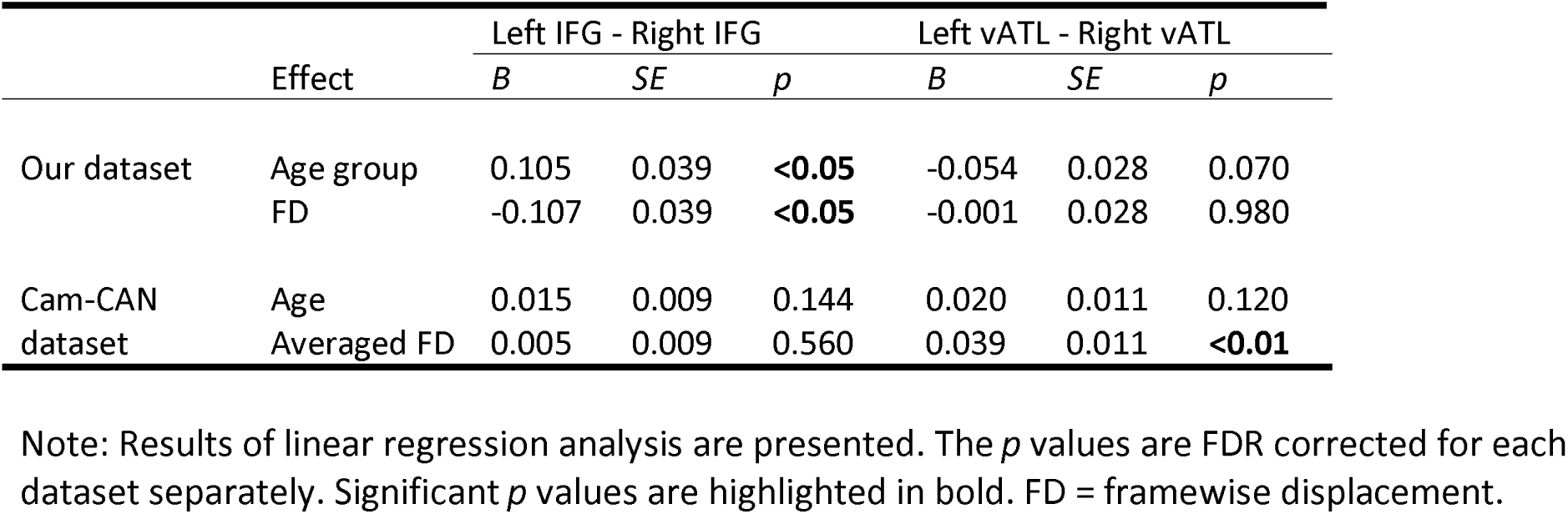
Results for age effects in the intrinsic functional connectivity analysis in two datasets.

**Table 2.** Results for linear regression testing the relationship between RSFC and expertise in our dataset.

| Effect |  | Left IFG - Right IFG |  |  | Left vATL - Right vATL |  |  |
| --- | --- | --- | --- | --- | --- | --- | --- |
|  |  | *B* | *SE* | *p* | *B* | *SE* | *p* |
| Semantic knowledge task | Age group | 0.161 | 0.041 | **<0.001** | -0.019 | 0.029 | 0.572 |
|  | Performance | -0.117 | 0.040 | **<0.01** | -0.094 | 0.028 | **<0.01** |
|  | FD | -0.163 | 0.042 | **<0.001** | -0.027 | 0.029 | 0.466 |
|  | Age group × Performance | -0.015 | 0.037 | 0.698 | -0.082 | 0.026 | **<0.01** |
| Semantic control task | Age group | 0.105 | 0.039 | **<0.05** | -0.054 | 0.028 | 0.147 |
|  | Performance | -0.016 | 0.035 | 0.968 | -0.005 | 0.025 | 0.968 |
|  | FD | -0.109 | 0.039 | **<0.05** | -0.001 | 0.028 | 0.968 |
|  | Age group × Performance | 0.050 | 0.035 | 0.303 | -0.004 | 0.025 | 0.968 |
| Non-semantic task | Age group | 0.112 | 0.039 | **<0.05** | -0.053 | 0.028 | 0.166 |
|  | Performance | -0.058 | 0.034 | 0.193 | -0.003 | 0.025 | 0.927 |
|  | FD | -0.121 | 0.040 | **<0.05** | -0.003 | 0.029 | 0.927 |
|  | Age group × Performance | -0.016 | 0.034 | 0.927 | 0.006 | 0.024 | 0.927 |

Note: The *p* values are FDR corrected for each task separately. Significant *p* values are highlighted in bold. FD = framewise displacement.

### Validation analyses

For the Cam-CAN dataset, we found no age effects on RSFC for left IFG -right IFG or left vATL - right vATL (Table 1 and Figure 2). In contrast, other IFG-vATL connections exhibited a significant decrease in strength with age (Supplementary Table S1 and Figure S1). For the task performance models, inclusion of STW scores improved the fit of the left vATL - right vATL model (*F* = 3.73, *p* < 0.05). As shown in Table 3 and Figure 4, people with better performance on the STW task exhibited weaker left vATL - right vATL connectivity (performance effect: B = -0.026, SE = 0.009, *p* < 0.05), replicating the effect of synonym judgement task in our dataset. This effect was present across ages (i.e., it did not interact with age). Unexpectedly, however, when controlling for STW performance, a positive effect of age emerged: connectivity between vATLs increased with age.

**Figure 4.**
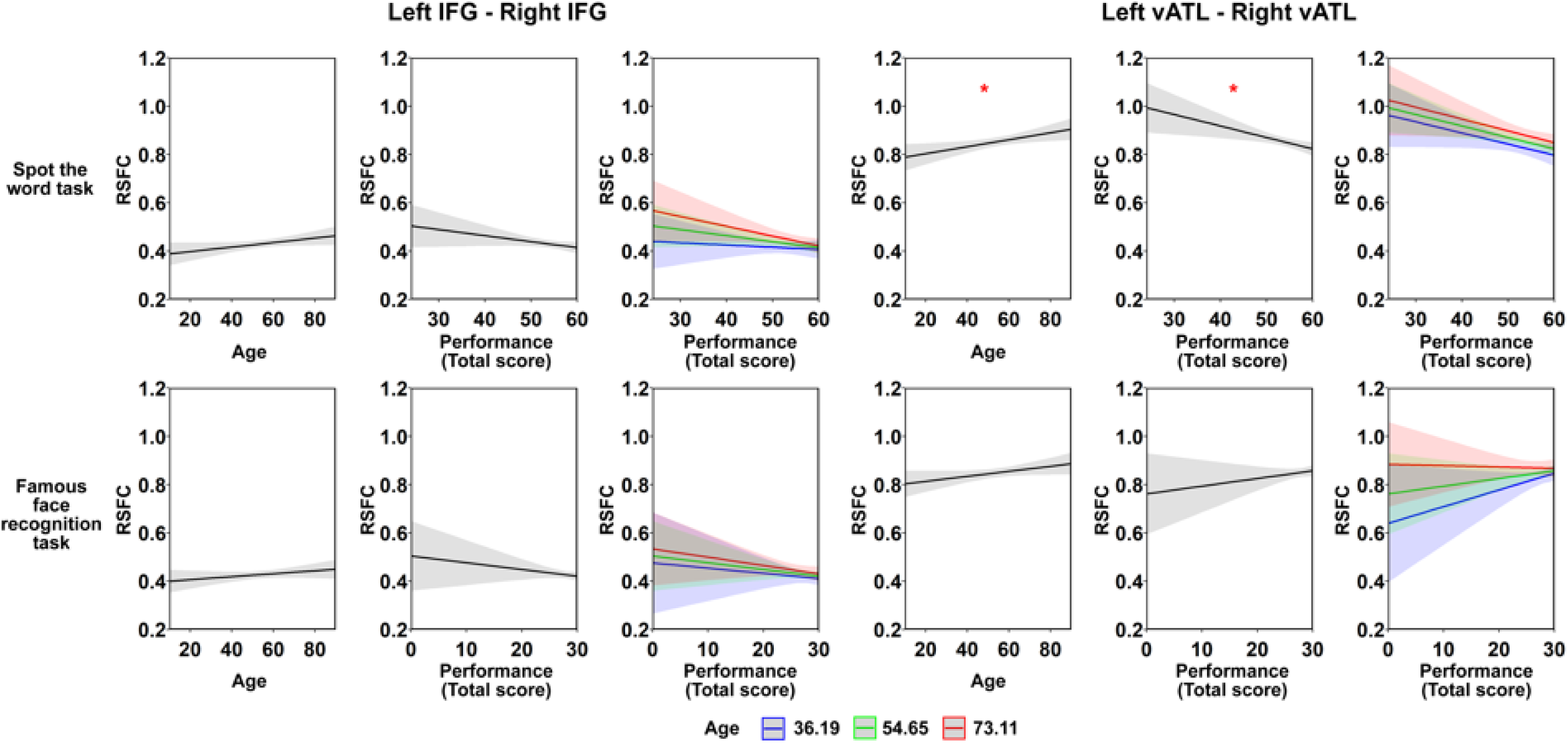
Results of linear regression model analysis for Cam-CAN dataset. This figure shows the modelled effects of age and task performance on RSFC between each pair of seed ROIs. Age × performance interaction is illustrated by plotting performance effects at the mean age and plus/minus 1 SD. Shadow areas and error bars indicate 95% confidence intervals. The asterisks indicate significance level after FDR correction within each task, * *p* < 0.05, ** *p* < 0.01, *** *p* < 0.001.

**Table 3.** Results for linear regression testing the relationship between RSFC and expertise in the Cam-CAN dataset.

|  |  | Left IFG - Right IFG |  |  | Left vATL - Right vATL |  |  |
| --- | --- | --- | --- | --- | --- | --- | --- |
| Effect |  | *B* | *SE* | *p* | *B* | *SE* | *p* |
| Spot the word task | Age | 0.017 | 0.009 | 0.129 | 0.027 | 0.011 | **<0.05** |
|  | Performance | -0.013 | 0.008 | 0.156 | -0.026 | 0.009 | **<0.05** |
|  | Averaged FD | 0.004 | 0.009 | 0.776 | 0.035 | 0.011 | **<0.01** |
|  | Age × Performance | -0.008 | 0.007 | 0.345 | -0.001 | 0.009 | 0.928 |
| Famous face recognition task | Age | 0.011 | 0.009 | 0.399 | 0.019 | 0.011 | 0.327 |
|  | Performance | -0.009 | 0.008 | 0.399 | 0.010 | 0.010 | 0.399 |
|  | Averaged FD | 0.006 | 0.009 | 0.611 | 0.039 | 0.011 | **<0.01** |
|  | Age × Performance | -0.002 | 0.007 | 0.760 | -0.012 | 0.008 | 0.335 |

Note: The *p* values are FDR corrected for each task separately. Significant *p* values are highlighted in bold. FD = framewise displacement.

Inclusion of STW scores did not improve the fit of the left IFG - right IFG model (*F* = 1.99, *p* = 0.14). This result diverges from our dataset, where we found that people with greater verbal semantic knowledge showed lower left IFG – right IFG connectivity. Here, the effect was in the same direction but was not statistically significant (two-tailed corrected *p* = 0.156). Inclusion of famous face performance did not improve fit of either left IFG – right IFG or left vATL – right vATL models (both *F* <= 1.39, both *p* >= 0.25), indicating that the observed effects were specific to verbal semantic knowledge. For the other connections (see Supplementary Table S3 and Figure S3), adding face recognition performance did improve model fit for the left IFG - right vATL RSFC (*F* = 3.64, *p* < 0.05). Better face recognition performance was linked with stronger correlation between these two regions (performance effect: B = 0.015, SE = 0.006, *p* < 0.05). STW performance did not predict the strength of any other connections.

## Discussion

The semantic network exhibits graded hemispheric specialisation, with verbal semantic processing more closely associated with left-hemisphere regions (Gainotti, 2014; Rice, Hoffman, et al., 2015; Rice, Lambon Ralph, et al., 2015; Snowden et al., 2012). We investigated the cognitive correlates of this specialisation using two independent resting-state fMRI datasets. Connectivity between left and right vATLs was lower in people with more advanced verbal semantic expertise, as indexed by tasks that probed their ability to recognise and understand low-frequency words. This effect was first found in our study of 86 young and older adults (Wu & Hoffman, 2023a) and then replicated in the Cam-CAN dataset (Shafto et al., 2014; Taylor et al., 2017), which includes over 600 participants across a wide range of ages. The effect was highly specific. Connectivity between vATLs was not correlated with performance on tasks that probed semantic control ability, non-semantic processing or face recognition. And while verbal semantic expertise was correlated with connectivity between left and right vATLs (and in the Wu & Hoffman data, between left and right IFGs), it was not correlated with the strength of within-hemisphere connections between semantic regions. Thus, there appears to be a specific and replicable relationship between the amount of verbal semantic knowledge a person has and the degree to which the left and right-hemisphere elements of their semantic system are functionally segregated from one another. This new insight is important for understanding the underlying causes of hemispheric specialisation in the semantic network.

Our interpretation of these findings is similar to that proposed by Behrmann, Plaut and colleagues for graded hemispheric specialisation in VOTC (Behrmann & Plaut, 2014; Behrmann & Plaut, 2020; Dundas et al., 2013, 2015; Plaut & Behrmann, 2011). It follows theories of graded semantic representation in the brain (Lambon Ralph et al., 2017; Plaut, 2002; Rice, Hoffman, et al., 2015) and involves three key assumptions. First, in line with the hub-and-spoke model (Lambon Ralph et al., 2017), we assume that the core computational function of the ATLs is to integrate inputs from a range of modality-specific processing streams. The ATLs generate semantic representations by extracting statistical regularities from these inputs. Second, we assume that the inputs received by different parts of the ATLs vary as a function of their connectivity with other neural systems. This variation occurs *within* each ATL, for example, the anterior portions of the superior middle temporal gyri are more strongly connected with auditory processing streams, while the anterior fusiform and parahippocampal gyri show stronger connections with VOTC (Binney et al., 2012). But more pertinently, we assume that it also occurs *across* hemispheres. Inter-hemispheric connections are much more abundant than cross-hemispheric connections (Suárez et al., 2018), resulting in a stronger connection between the left ATL and left-lateralized orthographic processing and language production regions. The final assumption is that specialisation in knowledge representation is determined by these variations in connectivity: neurons participate most strongly in representing concepts which are prominently featured in the inputs they receive. This specialisation is graded rather than absolute because representations are highly distributed across the entire ATL system (Schapiro et al., 2013).

Our results follow naturally from this account. The tests we used to measure verbal semantic expertise probe knowledge of low-frequency words that are typically acquired late in development, during adolescence and adulthood. As this knowledge is primarily acquired through exposure to written language (Cunningham & Stanovich, 1998) and written word recognition is left-lateralised (Dehaene et al., 2015), the left ATL is more able to represent this knowledge than the right. Thus, as people develop more sophisticated and diverse verbal semantic representations, left ATL regions become relatively specialised for this aspect of semantic processing, driving increased functional differentiation between left and right ATLs. This differentiation can be observed in terms of lower levels of resting-state connectivity between left and right ATLs in people whose verbal semantic representations are more developed.

Our account emphasises *written* word processing as a key driver of ATL specialisation. This claim is consistent with evidence that the left ATL bias for verbal semantics is strongest for written words and less prominent for spoken word processing (Hoffman & Lambon Ralph, 2018; Marinkovic et al., 2003; Rice, Lambon Ralph, et al., 2015; Spitsyna et al., 2006). It also aligns with another resting-state fMRI study which found that people with more verbal semantic expertise (measured using a written synonym judgement task) showed stronger connections between left ATL and orthographic cortex in left VOTC (Mollo et al., 2016). However, left ATL is also more connected with left-lateralised speech production systems than right ATL, and this connectivity bias might be an additional driver of specialisation for verbal semantic processing (Lambon Ralph et al., 2001; Schapiro et al., 2013).

If left ATL regions come to preferentially represent verbal semantic knowledge, are there other forms of semantic processing that rely preferentially on the right ATL? To answer this question, we need to consider which aspects of semantic representation rely on right-lateralised processing streams. Semantic representations for people is one candidate, given that face recognition processing in VOTC is right-lateralised (Behrmann & Plaut, 2020; Plaut & Behrmann, 2011). Patients with right ATL damage show more severe familiar face recognition deficits than those with left ATL damage (Rice et al., 2018; Snowden et al., 2012). If face/person expertise is the right-hemisphere counterpart to verbal semantic expertise, then we might expect people with highly developed person knowledge to also show a high level of segregation between ATLs. We did not find this: in the CamCAN data, we found no relationship between inter-ATL connectivity and performance on a famous face recognition task. It is important to note, however, in this test participants were asked name the face and to verbally provide their occupation and nationality. Although face processing may be right-lateralised, linking objects and people to their names is highly reliant on left ATL regions (Damasio et al., 2004; Lambon Ralph et al., 2001). Thus this task may have required a combination of representations with greater reliance on each ATL. Indeed, other than written words and faces, most meaningful stimuli seem to activate the ATLs in a relatively bilateral fashion (Hoffman & Lambon Ralph, 2018; Rice, Hoffman, et al., 2015) so we may not expect other knowledge types to correlate with functional segregation of the ATLs. For example, Alam et al. (2021) found that people who were highly proficient in identifying images of famous landmarks showed *higher* resting-state connectivity between left and right ATLs.

Although our main focus in this study was on left and right vATLs, we also tested whether resting-state connectivity between left and right IFGs correlates with semantic task performance. Here we found a more mixed picture. In the Wu and Hoffman (2023a) data, more advanced verbal semantic knowledge was associated with weaker inter-IFG connectivity, mirroring the result in the vATLs. In the CamCAN data, however, a smaller effect in the same direction was not statistically significant (two-tailed corrected *p* = 0.156). Further research is needed to establish the robustness of this effect. If, however, such an effect is found reliably, this may indicate the factors driving functional segregation in the ATLs have similar effects in the IFGs.

Functional connectivity did not correlate with performance on a feature-matching task designed to probe semantic control ability (Badre et al., 2005; Thompson-Schill et al., 1997). It is important to note that this task was not as exclusively verbal in nature as the semantic knowledge tasks. Although the stimuli were presented as written words, the semantic control task required participants to make judgements about specific object properties (colour, size) that are typically experienced through vision rather than language. This kind of non-verbal visual knowledge is less likely to show laterality effects, as object recognition is not subject to the same left-hemisphere biases as word recognition. In addition, the feature-matching task was not designed to probe the depth of participants’ semantic knowledge but rather their ability to resolve competition between active semantic representations. This semantic control ability relies heavily on the IFGs (Badre & Wagner, 2007; Hoffman et al., 2010; Thompson-Schill et al., 1997; Vitello & Rodd, 2015). While IFG activation for verbal semantic control processes is strongly left-lateralised, we have argued that right IFG also contributes to these processes when task demands are particularly high (Wu & Hoffman, 2023a). On this basis, one might expect people with poorer semantic control ability to show greater inter-IFG connectivity, as they rely more often on right IFG to support the functions of the left. This hypothesis would be best tested in future studies that manipulate the need for control in purely verbal semantic decisions (e.g., resolving competition between the meanings of ambiguous words).

Finally, we investigated how cross-hemispheric functional connectivity in the semantic system varies with age. In the Wu and Hoffman (2023a) data, correlations between left and right IFGs were higher in the older age group. This greater cross-hemispheric interaction could indicate that older people rely more on right IFG to support left IFG in regulating semantic activation (Hoffman & Morcom, 2018). This would be consistent with models that predict shifts towards greater bilaterality in later life to compensate for declines in cognitive function (Berlingeri et al., 2013; Cabeza, 2002). However, this effect was not replicated in the CamCAN data, which may indicate that it is not reliable or that it follows a non-linear trajectory in midlife which our analysis was not sensitive to. For the vATLs, the Wu and Hoffman (2023a) data showed a weak (non-significant) decrease in connection strength in older people that did not persist once verbal semantic knowledge was controlled for. In the CamCAN data, connectivity actually increased with age after controlling for verbal semantic expertise. These findings suggest that it is the acquisition of specific verbal knowledge that leads to increasing specialisation across ATLs, rather than simply the passage of time. Finally, in the Wu and Hoffman (2023a) data, the relationship between verbal semantic expertise and ATL segregation was only found in the older age group. The lack of an effect in the young age group may be due to a lack of variability in semantic knowledge in this group. Our younger participants were mostly aged in their early twenties and were almost all university students. It’s likely that levels of semantic knowledge varied less in this homogeneous group than they did in our older group. In contrast, the CamCAN data includes participants from a wide range of life stages. In this data, no interaction between the effect of verbal expertise and age was found, suggesting that the relationship between verbal semantic expertise and ATL segregation is stable across adulthood.

In summary, this study has established a specific and replicable relationship between individuals’ level of verbal semantic knowledge and the degree of functional segregation between their left and right vATLs. These findings support graded specialisation theories of ATL organisation (Guo et al., 2013; Lambon Ralph et al., 2017; Rice, Hoffman, et al., 2015; Rice, Lambon Ralph, et al., 2015). They suggest that this organisation is not innate but is rather an emergent consequence of developing expertise in verbal semantic knowledge. The underlying causes of this effect await confirmation in future work, though we propose that hemispheric biases in written word recognition processes are a likely driving factor.

## Acknowledgements

This work was supported by a BBSRC grant to P.H. (BB/T004444/1). Imaging was carried out at the Edinburgh Imaging Facility (www.ed.ac.uk/edinburgh-imaging), University of Edinburgh, which is part of the SINAPSE collaboration (www.sinapse.ac.uk). We are grateful to the University of Minnesota Center for Magnetic Resonance Research for sharing their neuroimaging sequences. We thank the Cambridge Centre for Ageing and Neuroscience (Cam-CAN) for collecting and sharing their data. Cam-CAN funding was provided by the UK Biotechnology and Biological Sciences Research Council (grant number BB/H008217/1), together with support from the UK Medical Research Council and University of Cambridge, UK. We would like to thank Dr. Haojie Wen for his valuable suggestions regarding resting-state data preprocessing procedures and Elizabeth Joyce, Junming Wei and Dr. Yueyang Zhang for their help with data collection and processing. We are also grateful to all research participants in this study. For the purpose of open access, the authors have applied a Creative Commons Attribution (CC BY) licence to any Author Accepted Manuscript version arising from this submission.

## CRediT authorship contribution statement

Wei Wu: Conceptualization, Investigation, Methodology, Formal analysis, Validation, Writing - original draft, Writing - review & editing, Visualization. Paul Hoffman: Conceptualization, Methodology, Validation, Writing - review & editing, Project administration, Supervision, Funding acquisition.

## Open practices

The current study uses data collected by Wu and Hoffman (2023). The behavioural data (https://doi.org/10.7488/ds/3845 and https://doi.org/10.7488/ds/3846), resting-state fMRI data and analysis code (https://osf.io/zbxt4) are publicly available. All task stimuli were obtained from the norms of Wu and Hoffman (2022), with digital study materials available at https://osf.io/9px7g.

