## Supplementary Information for "Verbal semantic expertise is associated with reduced functional connectivity between left and right anterior temporal lobes"

**Overview**

| Supplementary Table S1 | Results for age effects in the intrinsic functional connectivity analysis in two datasets |
| --- | --- |
| Supplementary Table S2 | Results for linear regression testing the relationship between RSFC and expertise in our dataset |
| Supplementary Table S3 | Results for linear regression testing the relationship between RSFC and expertise in the Cam-CAN dataset |
| Supplementary Figure S1 | Results of linear regression models for age effects |
| Supplementary Figure S2 | Results of linear regression models for our dataset |
| Supplementary Figure S3 | Results of linear regression models for Cam-CAN dataset |

**Supplementary Table S1. Results for age effects in the intrinsic functional connectivity analysis in two datasets**

|  |  | Left IFG - Left vATL | | | Left IFG - Right vATL | | | Right IFG - Left vATL | | | Right IFG - Right vATL | | |
| --- | --- | --- | --- | --- | --- | --- | --- | --- | --- | --- | --- | --- | --- |
|  | Effect | *B* | *SE* | *p* | *B* | *SE* | *p* | *B* | *SE* | *p* | *B* | *SE* | *p* |
| Our dataset | Age group | -0.002 | 0.024 | 0.99 | -0.0003 | 0.022 | 0.99 | -0.010 | 0.024 | 0.99 | -0.029 | 0.022 | 0.99 |
|  | FD | -0.022 | 0.025 | 0.99 | -0.005 | 0.023 | 0.99 | -0.004 | 0.025 | 0.99 | 0.004 | 0.023 | 0.99 |
| Cam-CAN dataset | Age | -0.034 | 0.006 | **<10^-6^** | -0.024 | 0.006 | **<10^-3^** | -0.041 | 0.007 | **<10^-7^** | -0.030 | 0.007 | **<10^-4^** |
|  | Averaged FD | 0.005 | 0.006 | 0.495 | 0.012 | 0.006 | 0.071 | 0.003 | 0.007 | 0.626 | 0.011 | 0.007 | 0.129 |

Note: Results of linear regression analysis (older contrasts young) are presented. The *p* values are FDR corrected for each dataset separately. Significant *p* values are highlighted in bold. FD = framewise displacement.

**Supplementary Table S2. Results for linear regression testing the relationship between RSFC and expertise in our dataset**

|  |  | Left IFG - Left vATL | | | Left IFG - Right vATL | | | Right IFG - Left vATL | | | Right IFG - Right vATL | | |
| --- | --- | --- | --- | --- | --- | --- | --- | --- | --- | --- | --- | --- | --- |
|  | Effect | *B* | *SE* | *p* | *B* | *SE* | *p* | *B* | *SE* | *p* | *B* | *SE* | *p* |
| Semantic knowledge task | Age group | 0.001 | 0.027 | 0.994 | 0.010 | 0.025 | 0.994 | -0.009 | 0.028 | 0.994 | -0.021 | 0.025 | 0.994 |
|  | Performance | -0.001 | 0.027 | 0.994 | -0.023 | 0.024 | 0.994 | -0.0002 | 0.027 | 0.994 | -0.014 | 0.025 | 0.994 |
|  | FD | -0.028 | 0.028 | 0.994 | -0.014 | 0.025 | 0.994 | -0.006 | 0.028 | 0.994 | -0.005 | 0.025 | 0.994 |
|  | Age group × Performance | 0.022 | 0.025 | 0.994 | -0.009 | 0.023 | 0.994 | 0.008 | 0.025 | 0.994 | 0.009 | 0.023 | 0.994 |
| Semantic control task | Age group | -0.002 | 0.024 | 0.983 | -0.0005 | 0.022 | 0.983 | -0.010 | 0.025 | 0.983 | -0.029 | 0.022 | 0.738 |
|  | Performance | 0.012 | 0.022 | 0.983 | 0.003 | 0.020 | 0.983 | 0.023 | 0.022 | 0.821 | 0.031 | 0.020 | 0.738 |
|  | FD | -0.021 | 0.025 | 0.902 | -0.005 | 0.023 | 0.983 | -0.002 | 0.025 | 0.983 | 0.008 | 0.022 | 0.983 |
|  | Age group × Performance | 0.024 | 0.022 | 0.821 | -0.027 | 0.020 | 0.738 | -0.001 | 0.022 | 0.983 | -0.041 | 0.020 | 0.646 |
| Non-semantic tasks | Age group | 0.001 | 0.025 | 0.981 | 0.001 | 0.023 | 0.981 | -0.007 | 0.025 | 0.981 | -0.029 | 0.023 | 0.981 |
|  | Performance | -0.020 | 0.022 | 0.981 | -0.012 | 0.020 | 0.981 | -0.026 | 0.022 | 0.981 | 0.009 | 0.020 | 0.981 |
|  | FD | -0.028 | 0.026 | 0.981 | -0.008 | 0.024 | 0.981 | -0.011 | 0.026 | 0.981 | 0.003 | 0.024 | 0.981 |
|  | Age group × Performance | -0.002 | 0.021 | 0.981 | -0.002 | 0.020 | 0.981 | -0.008 | 0.021 | 0.981 | 0.022 | 0.020 | 0.981 |

Note: The *p* values are FDR corrected for each task separately. Significant *p* values are highlighted in bold. FD = framewise displacement.

**Supplementary Table S3. Results for linear regression testing the relationship between RSFC and expertise in the Cam-CAN dataset**

|  |  | Left IFG - Left vATL | | | Left IFG - Right vATL | | | Right IFG - Left vATL | | | Right IFG - Right vATL | | |
| --- | --- | --- | --- | --- | --- | --- | --- | --- | --- | --- | --- | --- | --- |
|  | Effect | *B* | *SE* | *p* | *B* | *SE* | *p* | *B* | *SE* | *p* | *B* | *SE* | *p* |
| Spot the word task | Age | -0.037 | 0.006 | **<10^-7^** | -0.026 | 0.006 | **<10^-3^** | -0.041 | 0.007 | **<10^-7^** | -0.032 | 0.007 | **<10^-4^** |
|  | Performance | 0.010 | 0.005 | 0.167 | 0.007 | 0.005 | 0.370 | 0.003 | 0.006 | 0.757 | 0.008 | 0.006 | 0.370 |
|  | Averaged FD | 0.006 | 0.006 | 0.459 | 0.013 | 0.006 | 0.102 | 0.004 | 0.007 | 0.757 | 0.012 | 0.007 | 0.167 |
|  | Age × Performance | -0.002 | 0.005 | 0.757 | -0.002 | 0.005 | 0.778 | 0.007 | 0.006 | 0.370 | 0.001 | 0.006 | 0.927 |
| Famous face recognition task | Age | -0.031 | 0.006 | **<10^-4^** | -0.019 | 0.006 | **<0.05** | -0.038 | 0.007 | **<10^-5^** | -0.025 | 0.007 | **<0.01** |
|  | Performance | 0.009 | 0.006 | 0.208 | 0.015 | 0.006 | **<0.05** | 0.005 | 0.006 | 0.581 | 0.010 | 0.006 | 0.208 |
|  | Averaged FD | 0.004 | 0.006 | 0.581 | 0.012 | 0.006 | 0.134 | 0.003 | 0.007 | 0.726 | 0.011 | 0.007 | 0.208 |
|  | Age × Performance | 0.003 | 0.005 | 0.632 | 0.0005 | 0.005 | 0.918 | 0.006 | 0.005 | 0.326 | 0.006 | 0.005 | 0.326 |

Note: The *p* values are FDR corrected for each task separately. Significant *p* values are highlighted in bold. FD = framewise displacement.

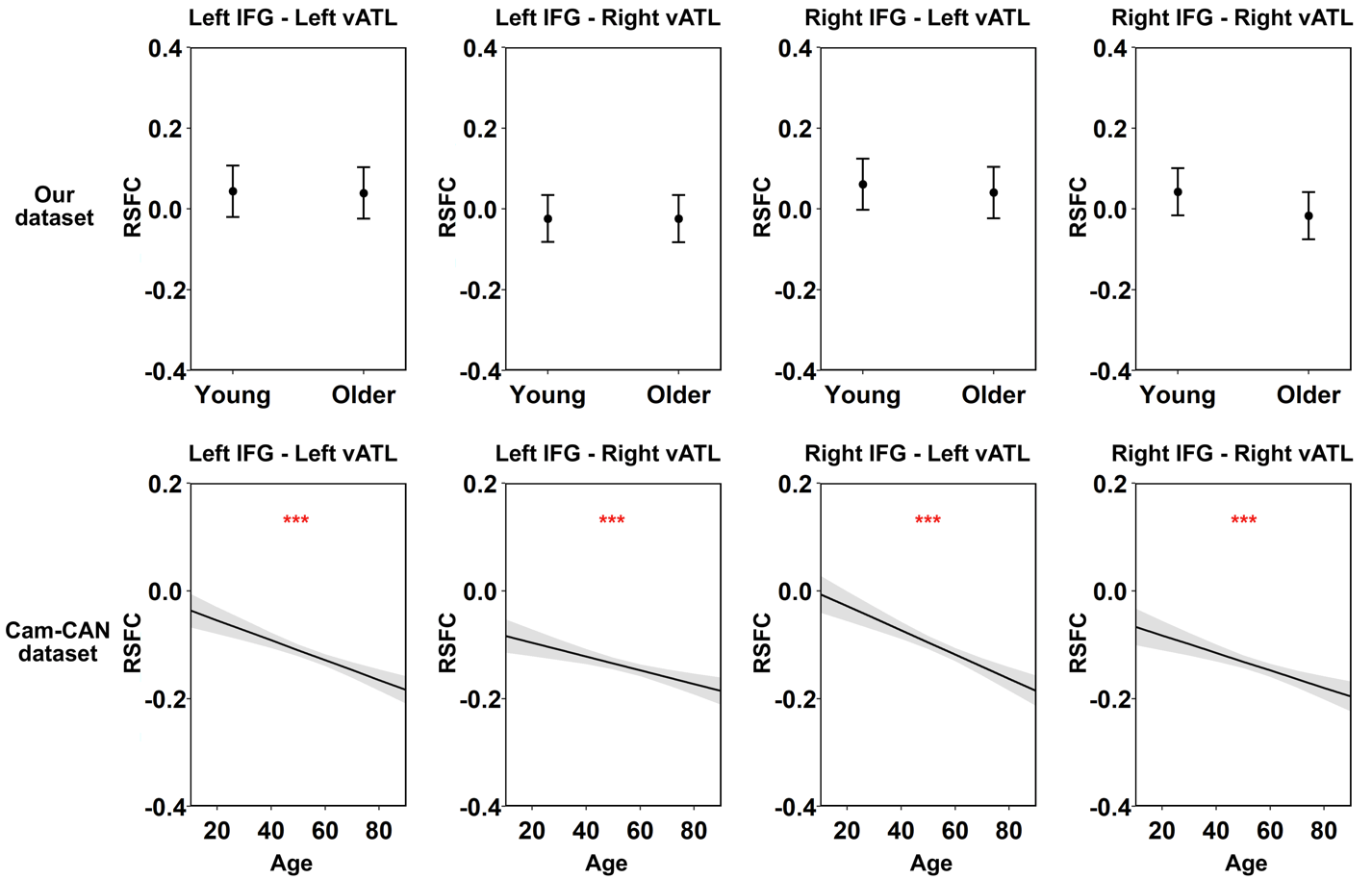

**Figure S1. Results of linear regression models for age effects. This figure shows the modelled effects of age on RSFC between each pair of seed ROIs in each task. Shadow areas and error bars indicate 95% confidence intervals. The asterisks indicate significance level after FDR correction within dataset, * *p* < 0.05, ** *p* < 0.01, *** *p* < 0.001.**

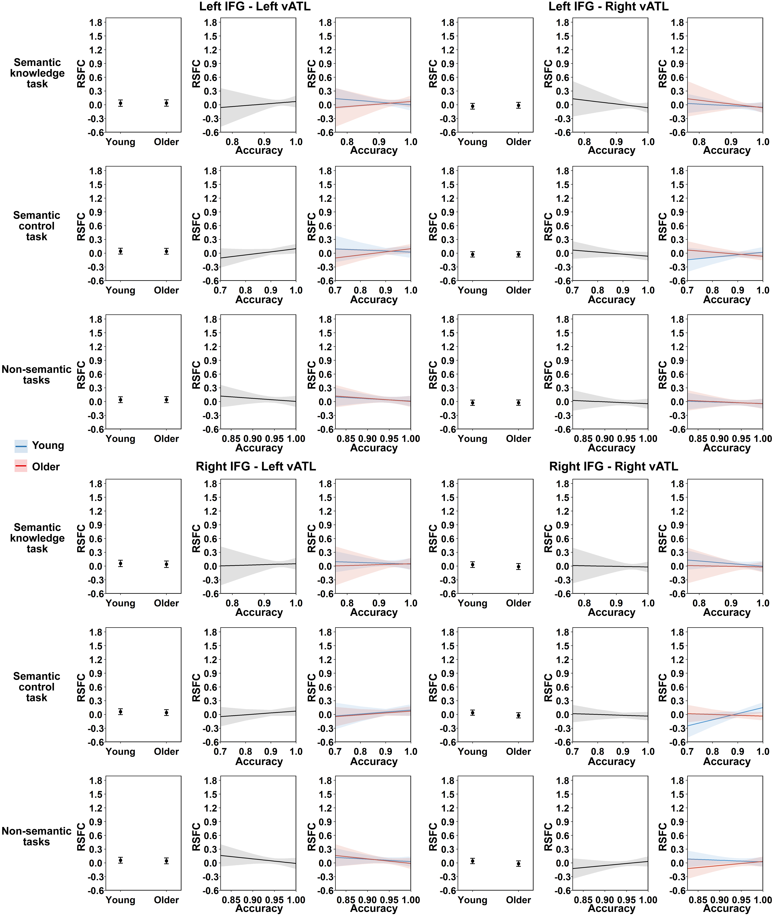

**Figure S2. Results of linear regression models for our dataset. This figure shows the modelled effects of age group and task performance on RSFC between each pair of seed ROIs. Shadow areas and error bars indicate 95% confidence intervals. The asterisks indicate significance level after FDR correction within each task, * *p* < 0.05, ** *p* < 0.01, *** *p* < 0.001.**

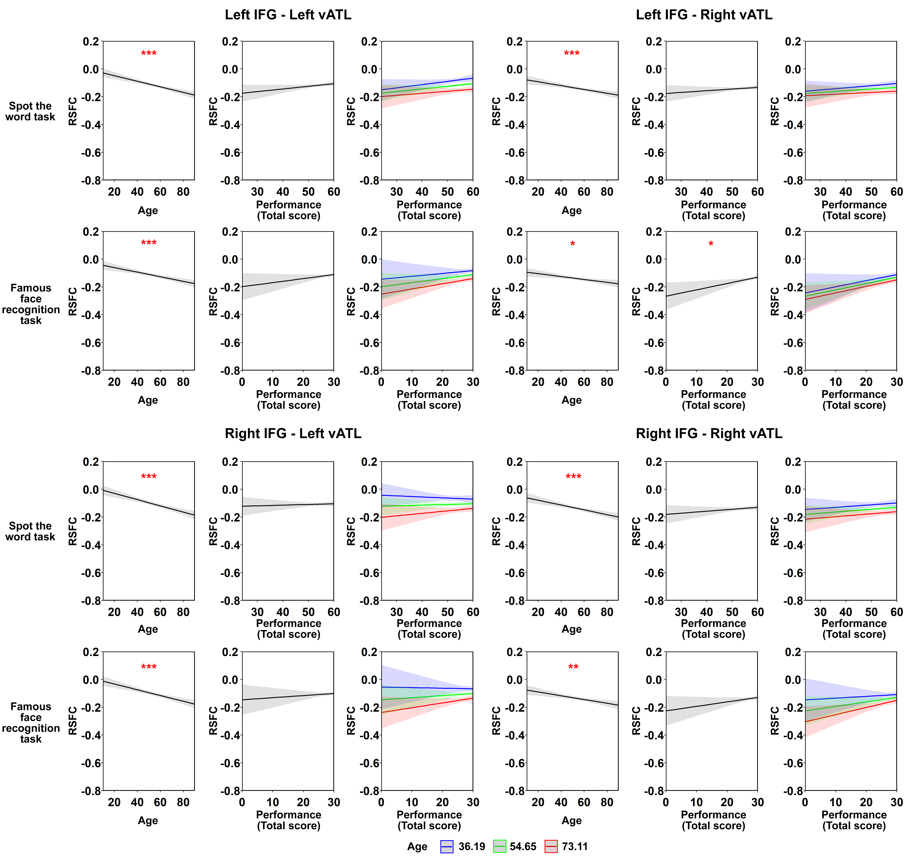

**Figure S3. Results of linear regression models for Cam-CAN dataset. This figure shows the modelled effects of age and task performance on RSFC between each pair of seed ROIs. Age × performance interaction is illustrated by plotting performance effects at the mean age and plus/minus 1 SD. Shadow areas and error bars indicate 95% confidence intervals. The asterisks indicate significance level after FDR correction within each task, * *p* < 0.05, ** *p* < 0.01, *** *p* < 0.001.**
